## Supplemental Information for "*Prm2* deficiency triggers a Reactive Oxygen Species (ROS)-mediated destruction cascade during epididymal sperm maturation in mice"

### Supplementary Information

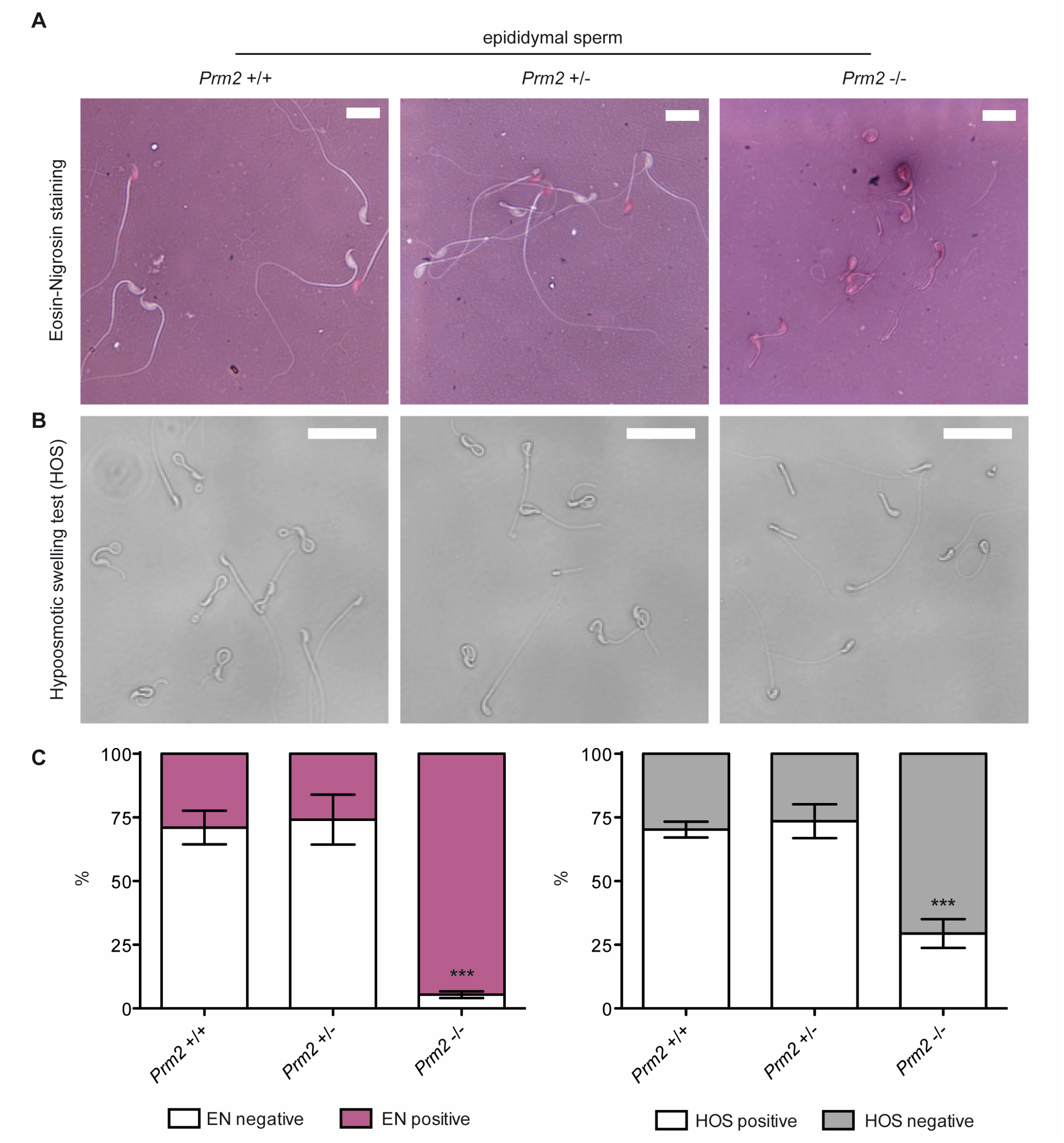


**Suppl. Fig. 1: Membrane integrity and sperm viability are decreased in *Prm2*-deficient mice.**
(**A**) Representative images of Eosin-Nigrosin (EN) stained epididymal sperm from *Prm2*$^{+/+}$, *Prm2*$^{+/-}$ and *Prm2*$^{-/-}$ males. Membrane damaged sperm stain pink, membrane intact sperm remain unstained. Scale bar: 10 µm. **(B)** Photomicrographs of epddiiymal sperm exposed to hypoosmotic conditions (HOS-test). Viable sperm display swelling and/or curling of the flagellum. Scale bar: 25 µm. **(C)** Quantification of EN staining and HOS-test. Bars represent mean values ±SD. Analysis comprised three biological replicates (n=3) per genotype. At least 200 sperm were evaluated per sample.

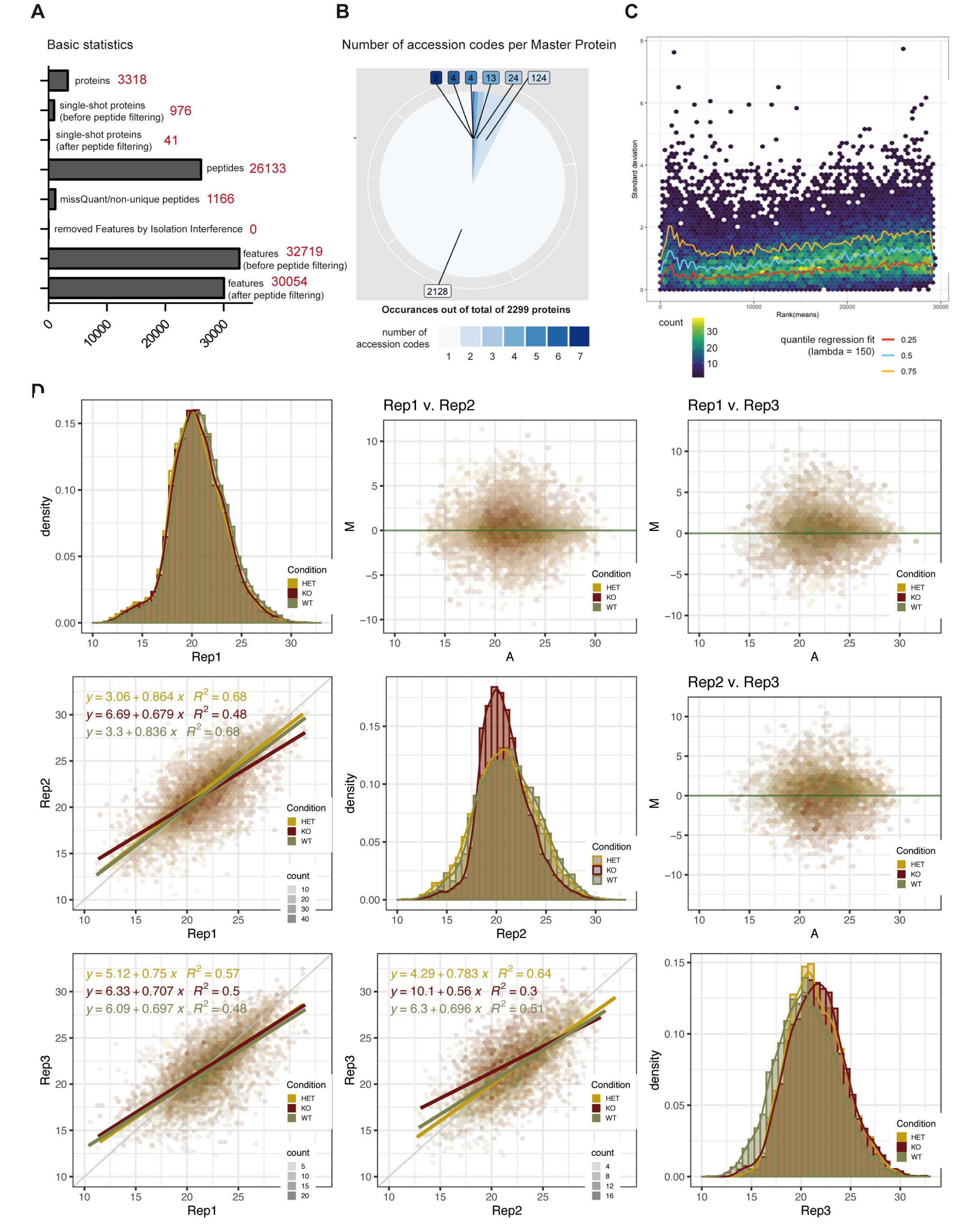


**Suppl. Fig. 2: Normalization and quality controls of MS data.**
**(A)** Basic statistics on the number of identified features, peptides and proteins (before and after peptide filtering). **(B)** Number of accession codes per master protein for 2299 proteins considered for label-free quantification. **(C)** Calibration of data using Variance Stabilizing Normalization (VSN) method. Data are $glog_2$ transformed to achieve variance stabilization. The hexogon plot depicts standard deviation versus rank of the mean. Count represents the number of data point falling into an hexagon. Lambda is an arbitrary factor to smooth quantile regression lines. **(D)** Normalization results for each replicate and condition are shown in diagonal. Scatter plots after normalization (bottom left) show a nice correlation among replicates. MA plots (upper right) display no systematic bias within the datasets.

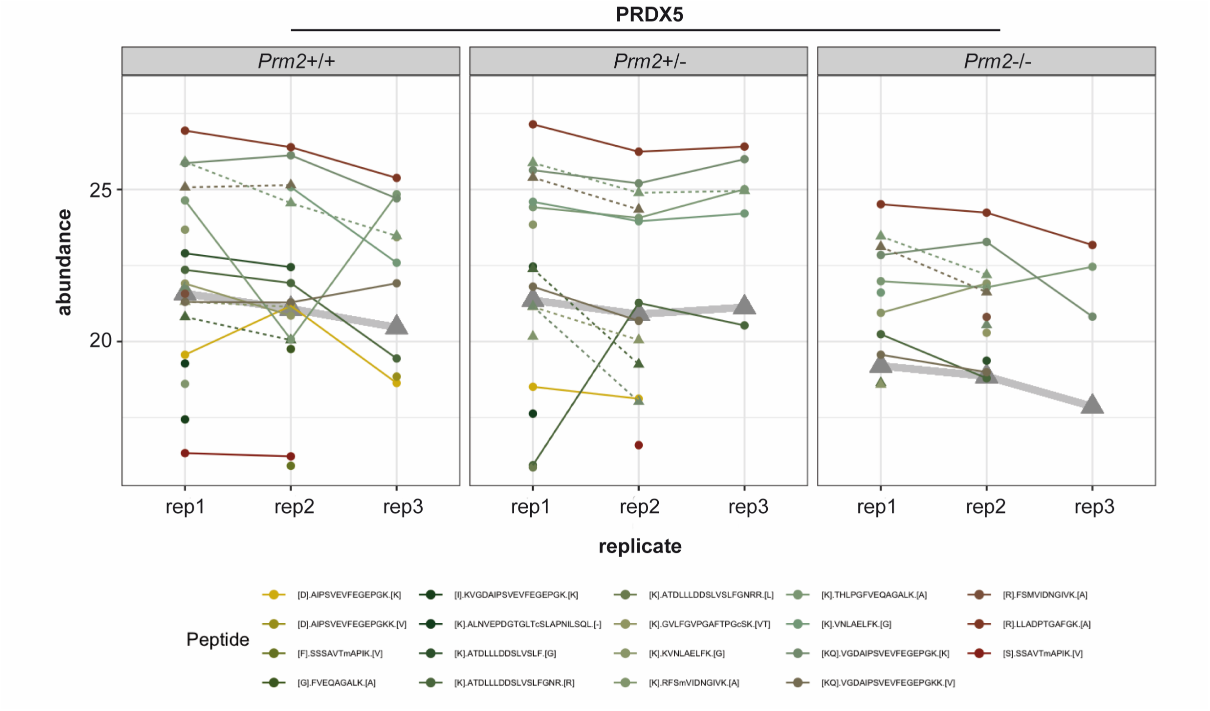


**Suppl. Fig. 3: Profile Plot PRDX5**

Relative abundance of identified PRDX5 peptides within each replicate of *Prm2*$^{+/+}$, *Prm2*$^{+/-}$ and *Prm2*$^{-/-}$ sperm retrieved from MS measurements by label-free quantification. Thickened grey lines resemble the abundance of the whole protein calculated by Tukey´s median polish.

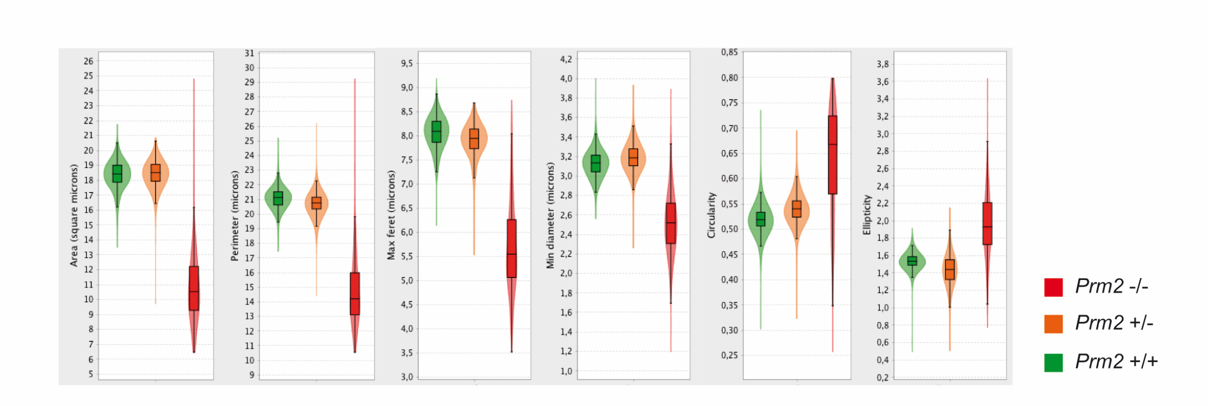


**Suppl. Fig. 4: *Prm2*-deficient epididymal sperm display a decreased head size.**
Violin plots displaying the head parameters area (µm$^2$), perimeter (µm), head length (µm), head width (µm), ellipticity and circularity. At least 420 individual sperm heads were evaluated per genotype. Plots are retrieved from the ImageJ plugin “Nuclear morphology analysis” (73).

**A**

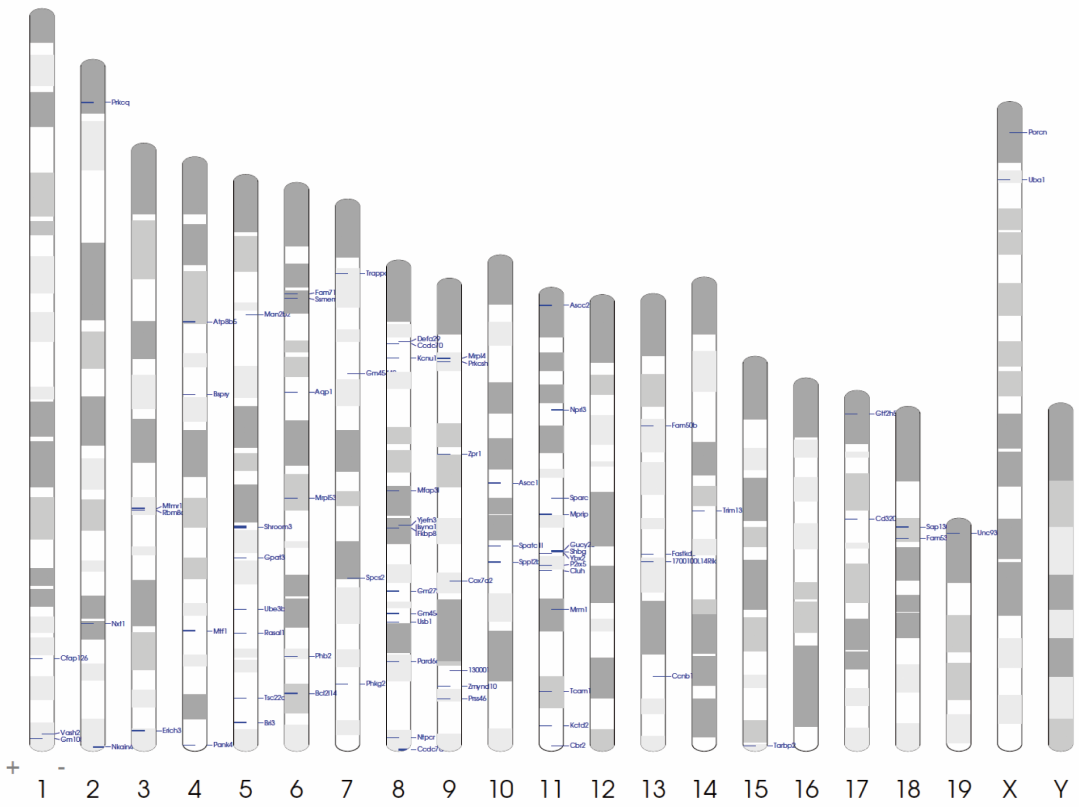


**B**

| GO biological process complete | Mus musculus REFLIST (22296) | upload (14) | expected | over/under | fold enrichment | raw p-value | FDR |
|---|---|---|---|---|---|---|---|
| protein modification by small protein conjugation (GO:0032446) | 462 | 5 | 0,29 | + | 17,24 | 0,000007 | 0,02 |
| protein modification by small protein conjugation or removal (GO:0070647) | 584 | 5 | 0,37 | + | 13,64 | 0,000021 | 0,04 |
| cellular protein modification process (GO:0006464) | 2268 | 9 | 1,42 | + | 6,32 | 0,000001 | 0,02 |
| protein modification process (GO:0036211) | 2268 | 9 | 1,42 | + | 6,32 | 0,000001 | 0,01 |
| macromolecule modification (GO:0043412) | 2441 | 9 | 1,53 | + | 5,87 | 0,000003 | 0,01 |
| cellular protein metabolic process (GO:0044267) | 2859 | 9 | 1,8 | + | 5,01 | 0,000010 | 0,03 |
| protein metabolic process (GO:0019538) | 3450 | 10 | 2,17 | + | 4,62 | 0,000004 | 0,02 |
| cellular macromolecule metabolic process (GO:0044260) | 3963 | 10 | 2,49 | + | 4,02 | 0,000016 | 0,04 |

**Suppl. Fig. 5: Analyses on DE genes identified by RNAseq of *Prm2*-deficient testes.**
**(A)** Idiogram showing the chromosomal localization of genes differentially upregulated in *Prm2*-deficient testes. Generated with Idiographica version 2.4 (www.rtools.cbrc.jp/idiographica/) (85). **(B)** Gene ontology analysis for enrichment of biological processes in the set of genes differentially downregulated in *Prm2*-deficient testes.
